# On the adaptive meaning of chemically insignificant cues in multitrophic caterpillar-ant-plant symbioses

**DOI:** 10.1101/2022.06.22.497176

**Authors:** Luan Dias Lima, Diego Santana Assis, Fábio Santos do Nascimento, Lucas Augusto Kaminski

**Author notes:** Author for correspondence: Lucas A. Kaminski.

## Abstract

Ants use cuticular hydrocarbons (CHCs) as chemical cues for recognition, which are exploited by myrmecophiles to usufruct the benefits of the social habit. We aimed to identify the functional role of CHCs of two riodinid caterpillar species that obligately associate with different multitrophic ant-plant symbioses. Experiments of ant acceptance of caterpillars showed the concerted action of larval-ant-organs that produce liquid rewards (tentacle nectary organs, TNOs) and luring signals (anterior tentacle organs, ATOs) was key to ant appeasement and larval survival. Experiments changing the symbiosis were often lethal for the caterpillars, mainly after emptying the secretions of their TNOs and their ATOs were never activated. Chemical profiles of caterpillars were insignificant. Field bioassays with chemically insignificant palatable insect prey attracted fewer ants, indicating that insignificant cues may reduce the probability of ant attacks. Thus, caterpillars control the emission of signals of ant-organs during symbiosis, whereas the cues of CHCs are silenced. It is possible that a trade-off exists between signals under control and non-controlled cues. Chemical insignificance may allow for a crucial period for caterpillars to make tactical decisions to appease aggressive ants and could be extended to other communication channels according to the sensorial universe of the receiver.

## 1. Background

Ants have existed since the Upper Cretaceous, approximately 100 million years ago, and constitute a dominant insect group in terms of abundance and biomass and are the main invertebrate predators in terrestrial ecosystems [1]. These insects, as with other eusocial insects, use intraspecific recognition systems and can distinguish between nestmates and non-nestmates using mainly chemical cues, namely compounds present in their cuticle called cuticular hydrocarbons (CHCs) (e.g. [2–4]). However, some organisms that interact with ants, i.e., myrmecophiles, can conceal information of chemical cues, allowing them to live with ants without being attacked and exploiting the benefits of the social habit, either protection, food, or shelter [1–2]. This intimacy and/or integration into ant nests and foraging territories on plants can be obtained through chemical strategies that include adaptive resemblance (such as mimicry, crypsis or masquerade) or not, such as chemical insignificance or conspicuousness [5–6].

Among adaptive resemblance strategies, chemical mimicry *sensu* [5] occurs when an organism is detected by a receiver (ant, in this case) as something interesting, resulting in action by the receiver. There are several examples of chemical mimicry strategy in the literature, mainly of social parasites that mimic the profiles of their host ants (e.g. [3,7–9]). These cases can be classified as aggressive mimicry (*sensu* [10]), since the emitter mimics the receiver to obtain benefits that directly harm the fitness of the receiver colony. Other strategies of adaptive resemblance include chemical crypsis, which occurs when the emitter organism is not detected due to the blending of its chemical cues with the background or chemical masquerade, when the organism is detected, but misidentified as something uninteresting, and thus not causing a reaction by the receiver [5]. These two kinds of strategies are broadly classified as chemical camouflage and have been reported for herbivorous insects as a way to reduce ant attacks (e.g. [6,11–14]).

Chemical insignificance occurs when an organism has few chemical cues due to the suppression or low concentration of cuticular compounds, which hampers its detection [5,15–16]. This strategy has been reported for some social parasitic ants and one caterpillar species (see [3,17]). Besides these strategies, ants can learn to recognise the CHC profiles of mutualistic partners, even when they are neither congruent with those of the ants nor the host plants, and thus avoid attacking them [18–20]. These non-congruent profiles may constitute conspicuousness strategy (*sensu* [6]).

Myrmecophilous butterflies (Lepidoptera: Lycaenidae and Riodinidae) live in association with ants during their immature stages (reviewed in [21]). In general, the ant- caterpillar interaction is trophobiotic, a kind of mutualistic interaction mediated by sugar-rich liquids that attract and serve as food resources for the ants, which in return offer protection against natural enemies or clean their partner caterpillars [1,20,22–23]. Such caterpillars use multimodal channels of communication to deal with ants, including tactile, chemical, and acoustic signalling, through morphological adaptations called ant- organs [22, 24–27]. Cues of CHCs can also mediate chemical strategies. Studies with the chemical profiles of myrmecophilous caterpillars with different kinds of interactions with ants (commensalism, mutualism, or parasitism), different degrees of specificity (facultative or obligate), and different feeding habits (herbivorous, omnivorous or carnivorous) have identified several kinds of chemical strategies (e.g. [6,17,28–29]). Thus, myrmecophilous caterpillars are excellent models for studying the ecology and evolution of different chemical strategies mediated by cuticular compounds [30].

Here, we assessed for the first time the chemical strategies mediated by CHCs of two caterpillar species of the family Riodinidae. Several species of this family engage in obligate trophobiotic interactions with ants (e.g. [31–32]), but the composition and function of the CHCs of these caterpillars are unknown. The studied species use similar host plants but obligately interact with different aggressive ant species that are phylogenetically distant (figure 1). Thus, the aim of this study was to perform chemical analyses and experiments in the field to identify the functional role of the CHC profiles of these caterpillars. Studies with species in the sister family Lycaenidae showed that trophobiotic caterpillars (that produce caloric rewards) generally have conspicuous CHCs that may appease ant aggressiveness or be associated with resource availability [6,20,33]. Based on this information, we hypothesised that the CHC profiles in the two caterpillar species would be informative to their respective tending ants and that, for this reason, they would be recognised as mutualistic partners and would not be attacked.

**Figure 1.**
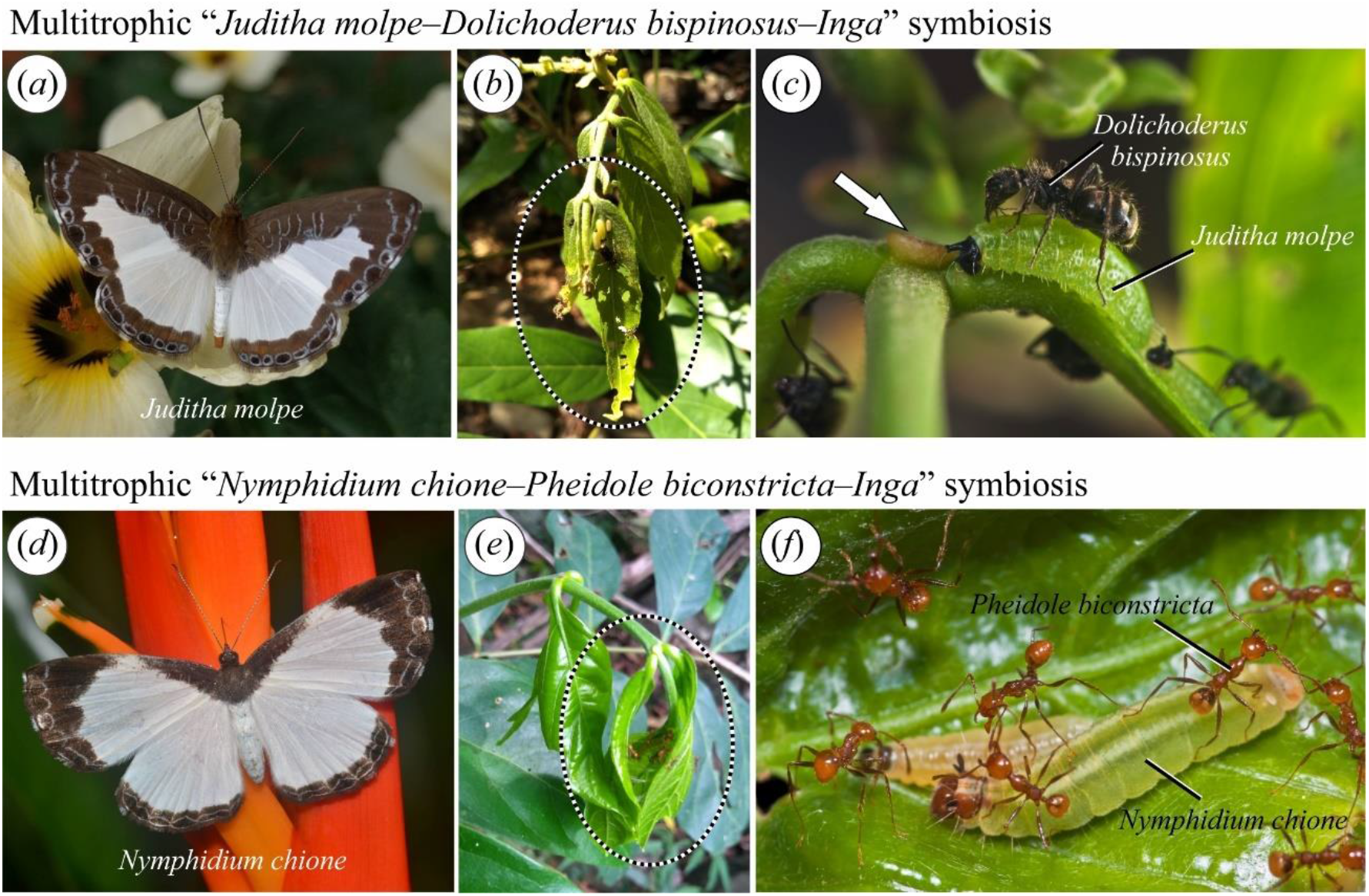
The studied multitrophic caterpillar-ant-plant symbioses: (a) *Juditha molpe* adult; (b) *Inga* branch with *J. molpe* caterpillars and *Dolichoderus bispinosus* ants (dashed ellipse); (c) detail of *J. molpe* caterpillars and *D. bispinosus* ants, note the *Inga* extrafloral nectary (white arrow); (d) *Nymphidium chione* adult; (e) *Inga* branch with *N. chione* caterpillars and *Pheidole biconstricta* ants (dashed ellipse); (f) detail of *N. chione* caterpillars and *P. biconstricta* ants. Photos by J. Movia^©^ (a), H. Soares Júnior^©^ (c,f), and by L.L. Mota^©^ (d).

## 2. Methods

### (a) Study site and organisms

The study was performed at the Serra das Araras Ecological Station (EESA) located in Porto Estrela (15º40’18” S; 57º13’47” W), state of Mato Grosso in the Center-West region of Brazil. The vegetation of EESA constitutes a transition between the Cerrado savanna and the Amazon forest [34–35]. The Amazon influence is stronger in the wetter areas of semideciduous forest and gallery forest and many caterpillar-ant-plant systems are typically Amazonian (e.g. [36]). Collections were carried out in the gallery forest of the Salobro river and Boca do José. Caterpillars were sampled in these sites by visually scanning vegetation. Sampling focused on two species of myrmecophilous butterflies abundant at EESA: *Juditha molpe* (Hübner, [1808]) and *Nymphidium chione* Bates, 1867 (Lepidoptera: Riodinidae). The caterpillars of these butterflies are classified as obligate myrmecophilous; *J. molpe* lives in association with *Dolichoderus bispinosus* (Olivier, 1792) (Dolichoderinae) and *N. chione* with *Pheidole biconstricta* Mayr, 1870 (Myrmicinae) (LAK, pers. observation) (figure 1). As observed for other obligate myrmecophilous riodinids, the caterpillars of *J. molpe* and *N. chione* are polyphagous throughout their geographic distribution, using mainly host plants bearing extrafloral nectaries [37–41]. The main host plants used by *J. molpe* and *N. chione* in areas of gallery forest at EESA are two species of *Inga* (Fabaceae). Plants of this genus have conspicuous extrafloral nectaries and attract many ants [42]. The ants, *D. bispinosus* and *P. biconstricta* (Myrmicinae), are top dominant ants in Neotropical forests and monopolise liquid food sources on vegetation [43–45]. In the study site, both species visit the extrafloral nectaries of *Inga* spp. but never occur together on the same plant (LAK, unpublished data).

Samples of the studied organisms were also collected for subsequent chemical analyses: *J. molpe* (first instar *n* = 6; second instar *n* = 8; third instar *n* = 3; fourth instar *n* = 2; last instar *n* = 1), *N. chione* (third instar *n* = 9; fourth instar *n* = 4; last instar *n* = 7), *D. bispinosus* (*n* = 100), *P. biconstricta* (*n* = 100), *Inga* sp. 1 (*n* = 10), and *Inga* sp. 2 (*n* = 10). These samples were frozen and kept at about −20°C until extraction of their cuticular compounds and subsequent chemical analyses.

### (b) Experiments of ant acceptance of caterpillars

Experiments were performed on ant acceptance of caterpillars to assess the role of ant- organs in ant-caterpillar interactions and the obligatoriness of this symbiosis. The experiments were performed on host plants in the field and consisted of two parts. For the control (without changing the associated ant), final instar (fourth or fifth) caterpillars of *J. molpe* and *N. chione* were removed from their host plants and transferred to another host plant of the same species occupied by their associated ant, i.e. *J. molpe* with *D. bispinosus* (*n* = 17) and *N. chione* with *P. biconstricta* (*n* = 15). For the treatment (change to the non-associated ant), the caterpillars were transferred to another host plant of the same species occupied by the non-associated ant species, i.e., *N. chione* with *D. bispinosus* (*n* = 15) and *J. molpe* with *P. biconstricta* (*n* = 16). All caterpillars were placed in small plastic pots with food, using careful manipulation with sterilized entomological tweezers, and being kept for 24 h without ants prior to carrying out the experiments. The caterpillars were then placed on plants in a manner that allowed the ants enough time to spontaneously find them. This practice was fundamental because quick movements and/or disturbances on the vegetation near the ants can immediately trigger aggressive responses from them, which would invalidate the acceptance trail.

The experimental plants (*Inga* spp.) were selected along a transect of approximately 1000 m based on plant species and presence of associated ants. Observations were made for 5 min, starting from the first contact between a worker ant and the placed caterpillar, which could occur after a variable amount of time. Behavioural responses were observed after these initial contacts and were classified at the end of the 5 min into two categories: 1) acceptance: ants tending the caterpillars without biting while antennating the ant-organs, i.e., tentacle nectary organs (TNOs), and anterior tentacle organs (ATOs) are constantly activated as described in the ‘enticement and binding’ process (*sensu* [25]); or 2) rejection: ants attacking the caterpillars with biting followed by recruitment of other ants. Each caterpillar and each plant were used only once in the experiments. Moreover, the action of ant-organs was observed *ad libitum* [46] during the acceptance experiments as well as their effects on the tending ants.

### (C) Chemical analyses

The cuticular chemical composition of the studied ants, caterpillars, and host plants was assessed by extracting the compounds following [6] by immersing the samples for 5 min in 5 ml of hexane (99%, SupraSolv, Merck, Germany), followed by removal from hexane with entomological tweezers that were previously cleaned with hexane. Pools of about 10 individuals (workers) per sample were used for tending ants, while for the caterpillars, one individual was used when it was in the last instar or a pool of up to five individuals when they were from first to the fourth instar. The cuticular compounds of the host plants were extracted following the same steps as for the insects, but with 20 ml of hexane. Each obtained extract was later evaporated under a fume hood and stored at −20°C until the moment of analyses when they were redissolved in hexane (95%, *n-*hexane, Macron, USA) and subsequently submitted to sonic baths.

All samples were analysed in a combined gas chromatography-mass spectrometer GC–MS (GC-QP2010 Plus, Shimadzu, Japan) equipped with a silica capillary column and using helium as a carrier gas following [47]. The conditions of the analyses were: injection performed manually with 2 µL using a glass syringe; splitless injection; oven temperature initially set at 150ºC increasing 3ºC/min until reaching 250°C, which was maintained for 10 min; injector temperature of 220ºC; temperatures of detectors and transfer lines were 250°C and 200°C, respectively; ionization energy of 70 eV and a range of 40–600 amu, with a scanning interval of 0.3 s.

Data were analysed with GC–MS software for Windows (Shimadzu Corporation), and the compounds were identified based on their mass spectra and with the aid of a standard solution (Sigma-Aldrich Chemical Co., USA) with synthetic linear hydrocarbons from C_21_ to C_40_, as well as by consulting the Wiley and NIST Libraries database. Compounds that were present in less than half of the individuals belonging to a group, and compounds contributing less than 0.5% to the total compounds, were excluded from the analyses.

### (d) Bioassays with chemically significant and insignificant model prey

Our initial hypothesis was that the cuticular compounds of the caterpillars would be informative (i.e. their own chemical signature) and, for this reason, they would be recognised and would not be attacked by the ant species they interact. However, the chemical analyses suggested that these caterpillars are chemically insignificant (see Results). Thus, we aimed to test the adaptive value of the chemical insignificance in the myrmecophily by performing field bioassays to examine the behavioural responses of the two ant species to the cuticular compounds of larvae of *Tenebrio molitor* Linnaeus, 1758 (Coleoptera: Tenebrionidae) a model of a palatable insect prey to be offered to the ants. We hypothesised that the chemically insignificant larvae would be less attractive and recognised as potential prey, reducing attack time and probability. The larvae used were previously killed by freezing at −20ºC and had a drop of hexane dripped on the cuticle (control) (chemically significant, with their normal CHCs) or that had their cuticular compounds extracted by hexane during 10 min at room temperature (treatment) (chemically insignificant, with reduction of CHCs). According to [48], the time of 10 min was sufficient to remove virtually all CHCs of *Blattella germanica* (Linnaeus, 1767) (Blattellidae) nymphs also working for Lepidoptera caterpillars [12].

The larvae were glued with white glue on leaves of removed branches of the *Inga* host plants of the studied caterpillars. Entomological tweezers, cleaned with hexane, were used to manipulate the larvae. The larvae were placed inside a small polystyrene foam cooler with rigid refrigerant bottles and kept cold until the moment of the bioassays. The removed branches with the experimental glued larvae were offered on the ground, at the base of the plants where ants were foraging, such that the ants could spontaneously find them. Pairs of control and treatment larvae were offered simultaneously, at a distance of at least 30 cm to avoid the recruitment effect of one larva on the other. A video of 5 min was recorded for each bioassay using a camera (Handycam Hdr-CX405 HD, Sony, Japan), starting from the first contact between a worker ant and the glued larva. The videos were later analysed and the total number of ants that touched a larva until the end of the bioassay was registered. Thirty replicates of this bioassay were performed (15 for each ant species).

### (e) Statistical analyses

The behavioural responses of the ants in the caterpillar acceptance experiments were assessed by performing the chi-square test (χ^2^) for comparison (or heterogeneity) between the proportions of the total number of caterpillars being accepted or rejected (control and treatment). The tested hypothesis was that the proportion of caterpillars accepted and rejected did not differ significantly between the control and treatment according to ant species.

The videos of the bioassays with chemically significant and chemically insignificant larvae were analysed by a Recruitment Index (RI) of the ants modified from [12]. The RI (mean ± SE) was calculated as the percentage of ants that touched larvae of the total number of ants recruited for the larvae (control and treatment) during 5 min. With the total number of ants recruited the RI was calculated as follows:

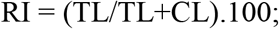

where RI is the Recruitment Index; TL is the number of ants that touched the treatment larva and; CL is the number of ants that touched the control larva. Thus, RIs values of 50 or near 50 indicate no difference in the recruitment between control and treatment, those higher than 50 indicate higher recruitment for treatment larva, and those lower than 50 indicate higher recruitment for control larva.

Moreover, the chi-square test (χ2) was also performed, as previously mentioned, for the bioassay videos of chemically significant larvae (with CHCs) and chemically insignificant larvae (with reduction of CHCs), but without subtracting 0.5 and comparing the total number of recruited ants for larvae (control and treatment). The hypothesis tested was that the proportion of ants recruited did not differ significantly between the control and treatment according to ant species.

We followed the methodology of [6] the percentages of absolute abundance of the cuticular compound extracts were calculated considering them as 100%. These data were then used to calculate the quantity of each separate compound (relative abundance) expressed as a percentage of the total occurrence of a class of substance. A principal component analysis (PCA) was performed using these data to verify similarities or differences among them. All analyses were performed using R environment [49].

## 3. Results

### (a) Experiments of ant acceptance of caterpillars

The caterpillar acceptance experiments (control) showed that *J. molpe* and *N. chione* are almost always accepted by their associated ants when transferred to another host plant (figures 1 and 2*a*), i.e., the caterpillars successfully appeased the ants. There was no heterogeneity regarding ant species, which means they had similar behavioural responses considering the caterpillars.

**Figure 2.**
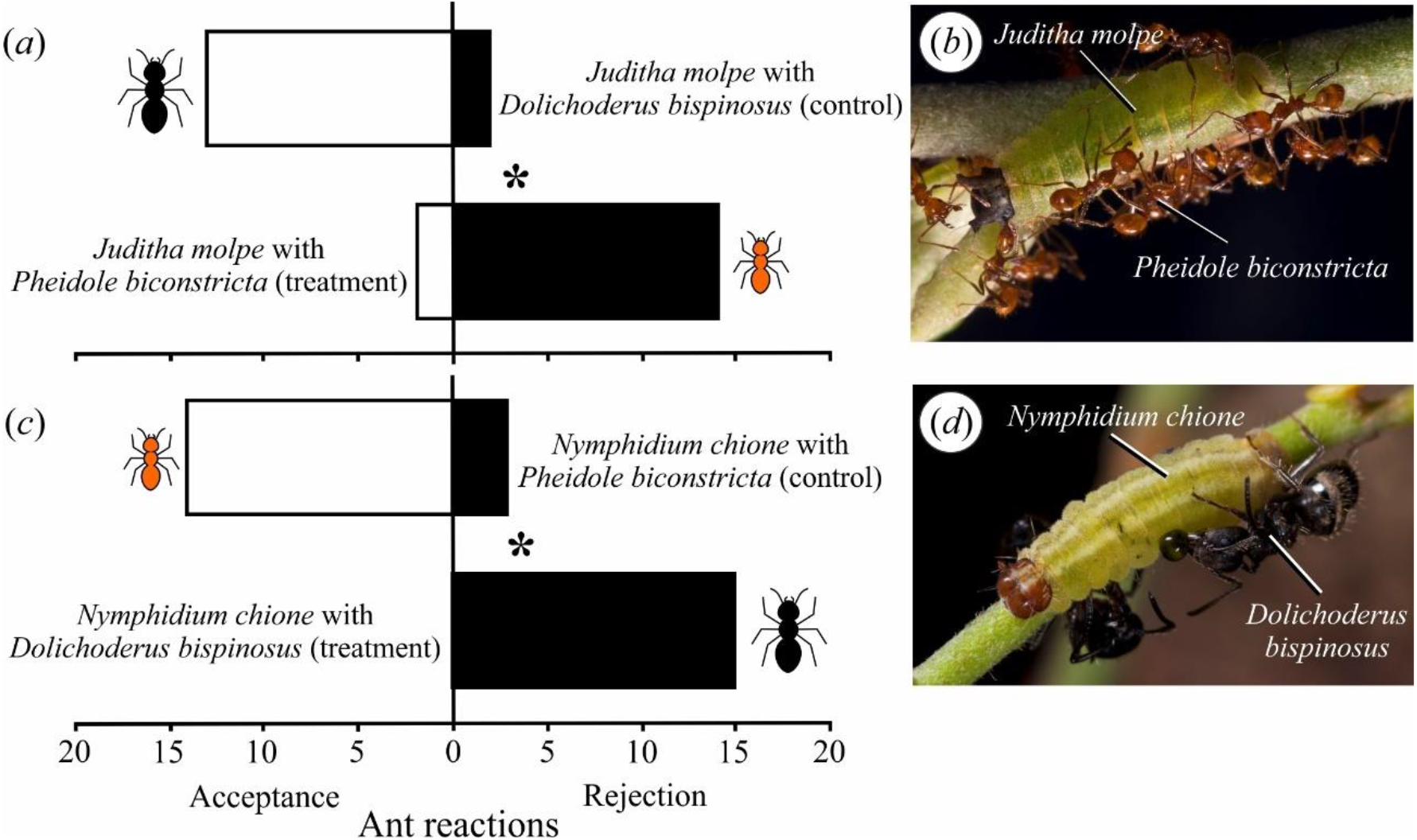
Ant reactions in ant acceptance of caterpillar experiments on *Inga* spp. (a) *Juditha molpe* caterpillars with associated tending ant *Dolichoderus bispinosus* (control) and with non-associated *Pheidole biconstricta* (treatment); (b) detail of *J. molpe* caterpillar attacked by *P. biconstricta* workers; (c) *Nymphidium chione* caterpillars with associated tending ant *P. biconstricta* (control) and with non-associated *D. bispinosus* (treatment); (d) detail of caterpillar attacked by *D. bispinosus* workers, note the hemolymph droplet leaking through the caterpillar tegument. (*) = significant values (χ^2^= 35.41; *df* = 1; *p* < 0.001). Photos by H. Soares Júnior^©^ (b,d).

The caterpillars used the concerted action of their ant-organs to appease the tending ants. The ants did not recognise the caterpillars immediately, only when they antennated the region near the ant-organs. The TNOs were promptly everted after rapid antennation (palpation), liberating a drop of secretion that was imbibed by the ants. After this first contact with the TNOs, ants started being recruited and the ATOs were everted. The action of this organ promoted excitement in the ants, which started “protecting” the caterpillar and reestablishing the symbiosis (figures 1*c,f*).

On the other hand, there was a significant difference between the aforementioned controls and treatments (χ^2^ = 35.41; *df* = 1; *p* < 0.001) for the experiments transferring caterpillars (treatment) between plants occupied with different ant species, which was often lethal for the two caterpillar species (figure 2). As in the control group, the caterpillars were firstly ignored until the ants found the TNOs. The, the antennating resulted in a drop of nectar being secreted that was imbibed by the ant. This appeasement lasted for a short amount of time, until the emptying of the secretions from the TNOs or when the ants antennated the region near the ATOs, which were never everted during the treatments. The emptying of the secretions of the TNOs seemed to result in more inspection of the caterpillar by the ants followed by biting. From this moment on, the ants started attacking the caterpillars and recruiting other ants (figures 2*b,d*). During the time that these experiments lasted, nearly all the caterpillars were killed. For *J. molpe* caterpillars with *P. biconstricta* (treatment), even soldiers with their robust mandibles, and who do not usually climb the plants, were recruited to help tear the caterpillars apart.

### (b) Chemical analyses

Samples of the caterpillars showed very low concentrations of CHCs, and so an extract was performed with all previously used samples totalling 20 caterpillars of each species to identify the compounds that remained in very low concentrations (electronic supplementary material, figure S1, table S1). The compounds identified in these extracts were mainly *n*-alkanes (C_25_–C_29_) for *J. molpe* and (C_21_–C_31_) for *N. chione* (electronic supplementary material, figure S1, table S1). The compounds identified in the ants were in a much greater quantity and concentration, with branched alkanes being the main compounds (electronic supplementary material, figure S1, table S1). There were significant differences between the cuticular compounds of the two species of caterpillars (*p* < 0.001) and also between those of caterpillars and ants (*p* < 0.001) (electronic supplementary material, figure S2). The compounds of the plants were not identified as they were visually much very and in a much greater quantity and concentration than those of the caterpillars, which shows that the caterpillars are not chemically camouflaged (electronic supplementary material, figure S1).

### (c) Bioassays with chemically significant and insignificant model prey

There were differences in the recruitment of *D. bispinosus* and *P. biconstricta* to *T. molitor* larvae between control and treatment with a tendency for more ant to recruitment to larvae with CHCs (control): 19.93 ± 3.92 for *D. bispinosus* on control and 10.6 ± 2.8 on treatment; 64.13 ± 15.87 for *P. biconstricta* on control and 50.73 ± 14.26 on treatment (χ^2^ = 53.31; *df* = 13; *p* < 0.001) (figure 3).

**Figure 3.**
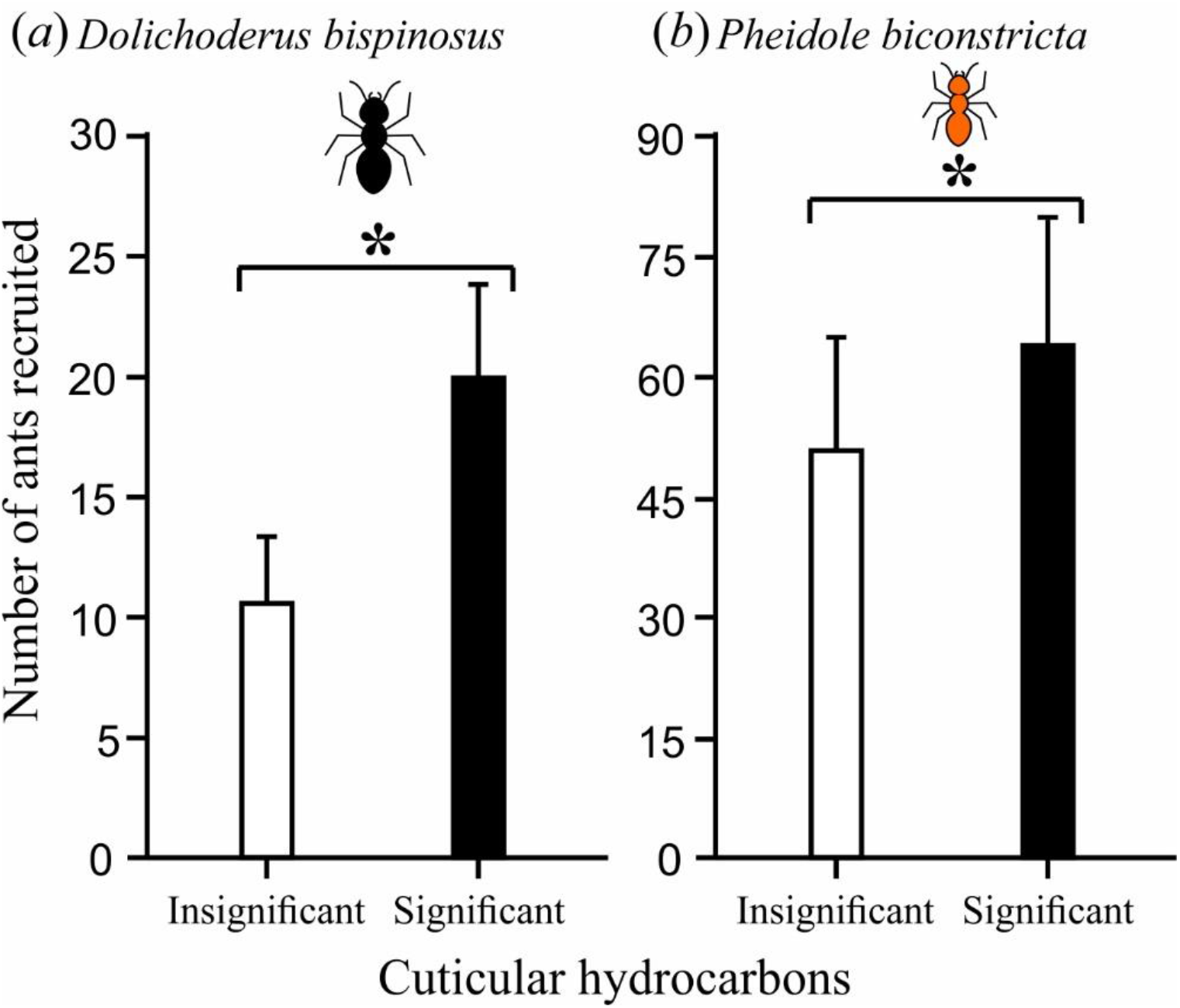
Number of ants recruited in bioassays with palatable model prey (larvae of *Tenebrio molitor*): chemically insignificant (with reduction of cuticular hydrocarbons, white bars) and chemically significant (with cuticular hydrocarbons, black bars). (a) *Dolichoderus bispinosus*; (b) *Pheidole biconstricta*. Values are means ± SE; (*) = significant values (χ^2^ = 53.31; *df* = 13; *p* < 0.001).

However, there was heterogeneity regarding ant species (χ^2^ = 406.76; *df* = 13; *p* <0.001). There was higher recruitment of *D. bispinosus* for larvae of *T. molitor* with CHCs than to larvae with reduced CHCs (RI = 35.21 ± 6.82) and there was a clear tendency for higher recruitment to larvae with CHCs (χ^2^ = 42.79; *df* = 14; *p* < 0.001) (figure 3*a*).

The recruitment of *P. biconstricta* was also higher for larvae with CHCs than those with reduced CHCs (RI = 43.45 ± 5.95), however, there was not a clear tendency for higher recruitment to larvae with CHCs (figure 3*b*).

## 4. Discussion

The studied caterpillar species were generally accepted by their associated tending ants and killed when they were transferred to host plants occupied by non-associated ants. Experiments changing myrmecophilous riodinid caterpillars from their associated ants have demonstrated that they may be accepted by some ant species, but sometimes attacked by others [31,50–51]. In these studies, failure in signalling has been proposed as a factor that could trigger the observed negative reactions, a hypothesis that is corroborated by the experiments of the present study as the caterpillars controlled the emission of specific signals from their ant-organs. There is evidence that ants use CHCs to recognise nestmates and non-nestmates and attack species with which they do not have mutualistic interactions (e.g. [1,15,52]). It has been shown that some CHCs of caterpillars associated with many ant species (facultative myrmecophily) may trigger aggressive responses or appeasement in different ant species [8].

The caterpillars studied here are presumed to have mutualistic interactions with ants, so our initial hypothesis that caterpillars would have informative CHC profiles, as reported for other trophobiotic species (e.g. [6,18–20]), was not confirmed. The strategies of chemical mimicry and chemical camouflage were discarded due to the low concentrations and number of compounds and incongruence with the CHC profiles of the ants or host plants. Moreover, the present study demonstrates, through chemical analyses, that the caterpillars use the chemical insignificance strategy, thus silencing their cuticular chemical cues. Chemical insignificance has been reported as an effective strategy to decrease recognition and attacks by ants [3,5,15]. This strategy has been commonly reported for several ant parasite taxa, i.e., ants, arachnids, isopods, collembolans, coleopterans, dipterans, gastropods, and zygentomans [3 and references therein,8,53–55]. Within Lepidoptera, this strategy has already been reported for the pupa of *Feniseca tarquinius* (Fabricius, 1793) [28] and the caterpillar of *Arhopala zylda* Corbet, 1941 [17]. However, the adaptive basis of this chemical insignificance has never been tested and continues to be a little-understood phenomenon among myrmecophilous organisms.

In the present study, caterpillars were observed using their ant-organs when discovered by ants. Glands that produce caloric rewards (TNOs) were active regardless of the tending ant species. However, other factors, such as luring signals from caterpillar ATOs, may have influenced associated ants not to attack during the trials but were not used with non-associated ants. The concerted use of ant-organs was also observed for the riodinid caterpillar *Thisbe irenea* (Stoll, 1780), which could appease associated ants yet never evert ATOs when attacked [25].

The myrmecophilous riodinid species of the subtribes Lemoniadina and Nymphidiina (*sensu* [56]) have TNOs that secrete nutritional substances and ATOs that are everted during symbiotic interactions with ants [25,57–59]. According to [17], the caterpillars of *Arhopala* (Lycaenidae) generally have dorsal nectary organs (DNOs) that produce nectar rewards analogous to the TNOs in riodinid caterpillars, posterior tentacle organs (TOs) analogous to ATOs in riodinid caterpillars [51], and pore cupola organs (PCOs) that supposedly produce substances that decrease attacks by the ant *Crematogaster decamera* Forel, 1910. Indeed, other authors have shown that DNO secretions from lycaenids can appease ants [60].

The chemical insignificance strategy has arisen independently in the two butterfly families of species with trophobiotic caterpillars on plants, as well as in other myrmecophiles with social parasitic lifestyles [3,17,28]. In addition to chemical analysis, the results of the field bioassays show that the removal of CHCs decreases the recruitment of ants to a palatable model insect, which suggests a possible adaptive value of the chemical insignificance in this case by allowing a crucial period of time for the caterpillar to decide on tactics to use to appease aggressive ants. This can be added to the fact that ants will even prey on mutualistic aphids when they do not secrete sugar-rich liquids (e.g. [61–62]). In the present study, caterpillars that were transferred to another host plant were attacked and killed by non-associated tending ants. This suggests that other factors must be acting in the establishment of stable caterpillar-ant symbioses, but reinforces the importance of a chemical strategy for caterpillars to keep close to aggressive ants and avoid recognition as prey, thereby reducing the possibility of attacks. Caterpillars of the present study were also observed using secretions as a way to keep close and appease ants, as well as avoid attacks, which worked only with their associated ant partner. Something similar was observed by [17], when *Arhopala* caterpillars were transferred to other plants, although different for *A. zylda* which also used the chemical insignificance strategy and was ignored by ants. The riodinids studied here may use the same tactics as *Arhopala* caterpillars as they also have ant-organs and secrete nectar rewards for aggressive associated ants to avoid attacks.

According to [25] the ATO chemistry is important for caterpillars to be accepted by certain species of ants and it does not function with others that kill them. This could be related to chemical signals that are specific to their associated ants and do not work with ants from different subfamilies as they may have coevolved. This is because there are chemical cues that can provide information unintentionally and can act directly as precursors for the evolution of chemical signals that evolved due to a specific effect, providing information intentionally, and work because the receiver’s response also evolved [63–64]. Thus, understanding the difference between cues and signals is essential to understanding the role of CHCs in myrmecophilous systems as cues and signals can act simultaneously [64] and can interfere with the fitness of myrmecophiles.

The present study found an expression of chemical signals from ant-organs that are intentional, however, the cuticular chemical cues do not allow this expression. The results reveal a possible trade-off between controlled signals and non-controlled cues in obligate multitrophic caterpillar-ant-plant symbioses. In this context, the insignificance of chemical cues may be adaptive in multitrophic interactions that involve complex multimodal signals by avoiding deaths in unpleasant encounters with non-associated ants.

The insignificance of chemical cues is a strategy that could be extended to other channels of communication according to the sensorial universe of the receiver and the degree of intimacy of the interaction. Caterpillars use a multimodal system to deal with ants [27], but this was not evaluated in the present study. For example, the apparent loss of the ability to produce sounds by caterpillars of some species of *Nymphidium* (see [26]) may be related to selection for cue insignificance. Another remaining question is whether chemical insignificance can also decrease attacks of other natural enemies such as chemically oriented predators and parasitoids.

## Ethics

The collection and transport of the specimens were authorised by Sistema de Autorização e Informação em Biodiversidade (SISBIO) (licence no. 62345-1).

## Data Accessibility

The data that support the findings of this study are openly available on Figshare [65]. Supplementary information is provided in the electronic supplementary material [66].

## Authors’ contributions

L.D.L.: conceptualization, data curation, formal analysis, funding acquisition; investigation; methodology; software; validation; visualization; writing – original draft; writing – review & editing; D.S.A.: data curation; formal analysis; software; validation; visualization; writing – review & editing; F.S.N.: funding acquisition; investigation; project administration; resources; supervision; validation; visualization; writing – review & editing; L.A.K.: conceptualization; data curation; funding acquisition; investigation; methodology; project administration; resources; supervision; validation; visualization; writing – original draft; writing – review & editing. All authors gave final approval for publication and agree to be held accountable for the work performed therein.

## Acknowledgments

We are grateful to: Marcelo L. F. Andrade (ESEC Serra das Araras) for permission and logistic support during fieldwork; Izabel C. C. Turatti for the assistance with identification of chemical compounds; Helena P. Romanowski for help with statistical analyses; Ana Paula Goetz for assistance in the field; Joaquim Movia and Luísa L. Mota for the butterfly photos; Hélio S. Júnior for the beautiful caterpillar-ant macrophotographs; and Erik R. Wild for revising the English text. We especially thank Thiago J. Izzo and students Brayan R. Massaroli, Carolina Nardes, Kamila P. Thomas, Renata S. Souza and Thais Conceição of the course Ecologia de Campo (2017) of the PPG-Ecologia e Conservação da Biodiversidade of the Universidade Federal de Mato Grosso for the opportunity to get to know the ecological station and assistance with the acceptance experiments.

