## Supplementary figure 1 for "On the adaptive meaning of chemically insignificant cues in multitrophic caterpillar-ant-plant symbioses"

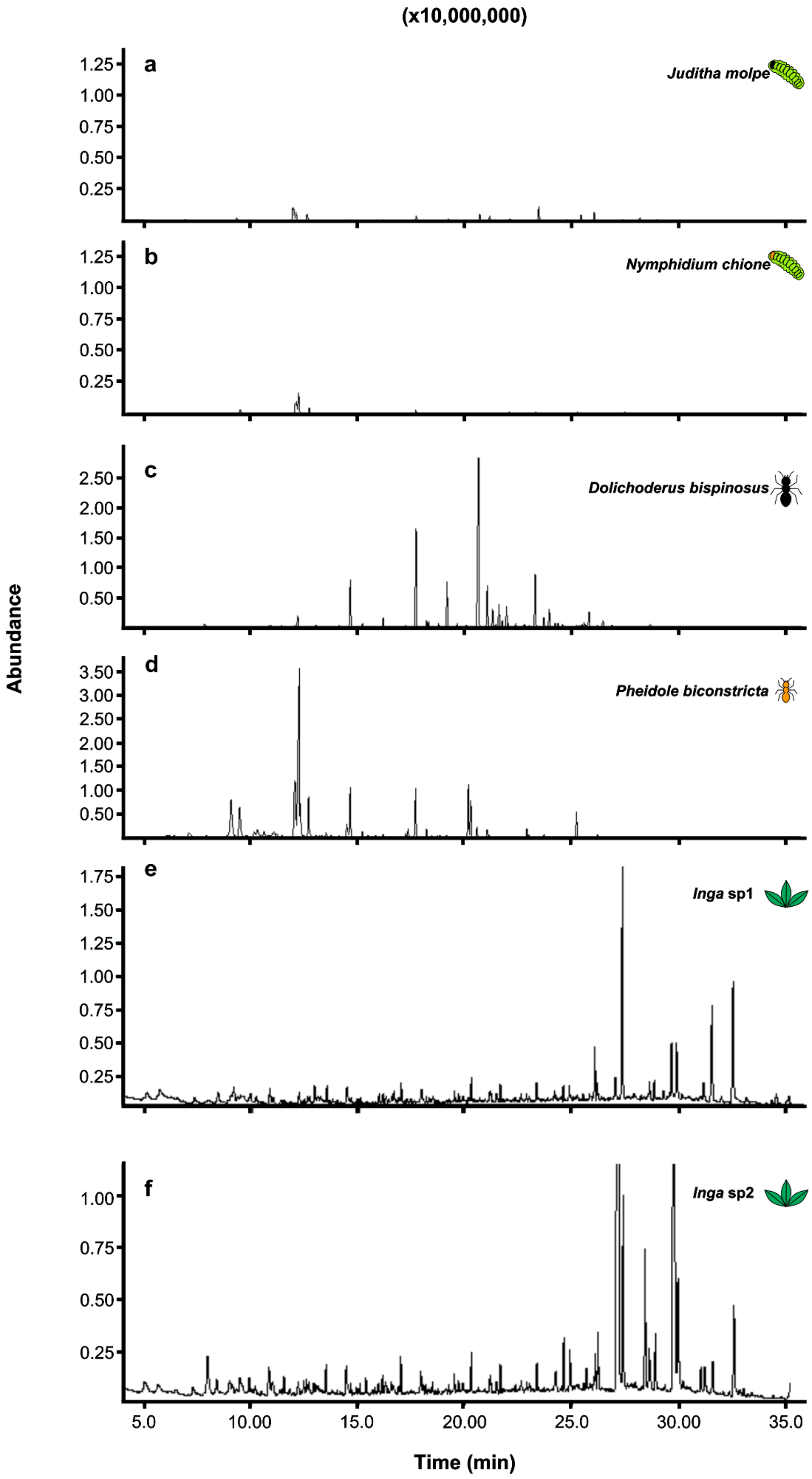


**Figure S1.** Chromatograms of the cuticular compounds. (a) *Nymphidium chione* caterpillars; (b) *Juditha molpe* caterpillars; (c) *Dolichoderus bispinosus* workers; (d) *Pheidole biconstricta* workers; (e) *Inga* sp. 1 leaves and f) *Inga* sp. 2 leaves.
