## Supplementary figure 2 for "On the adaptive meaning of chemically insignificant cues in multitrophic caterpillar-ant-plant symbioses"

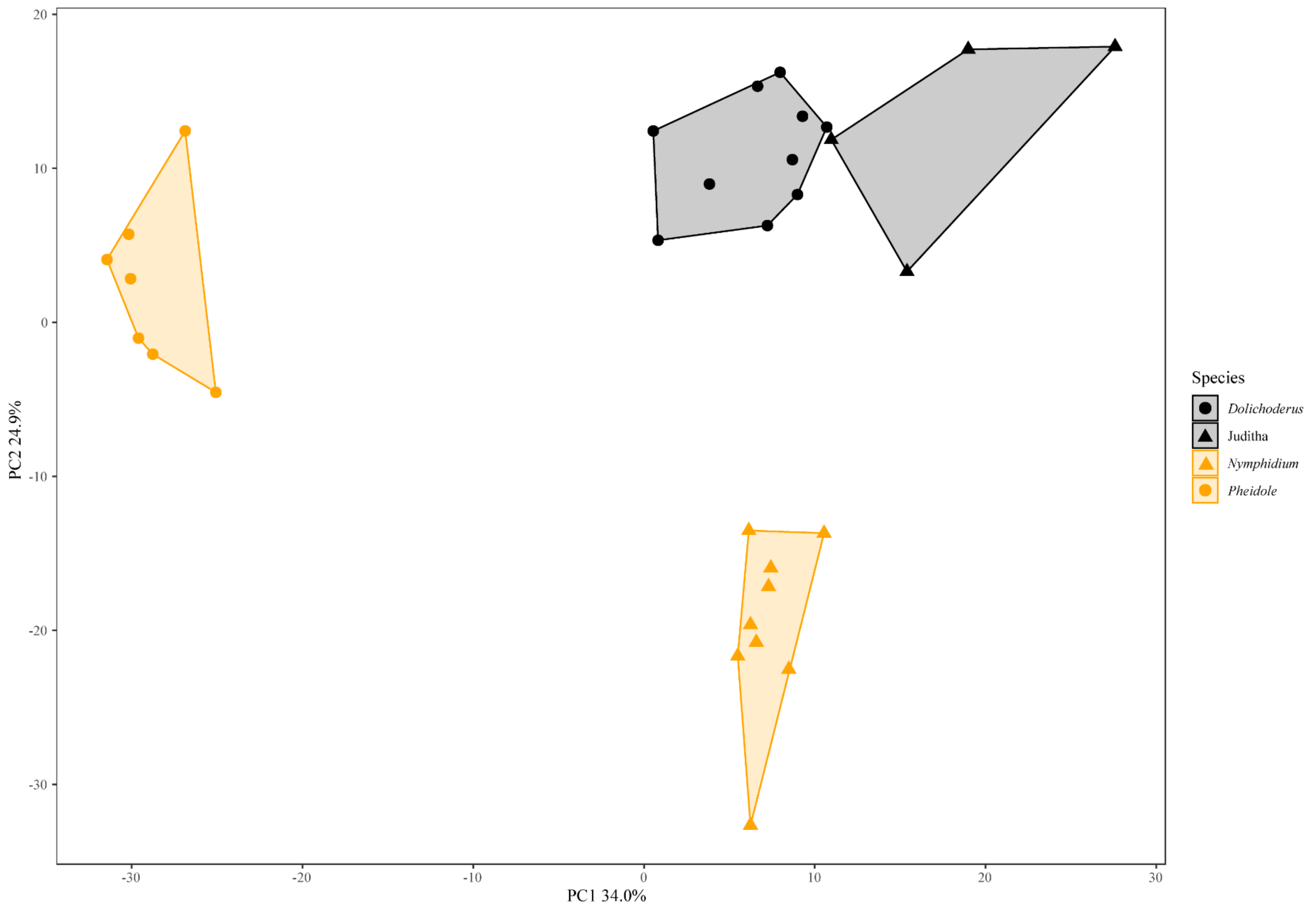


**Figure S2.** Principal component analysis (PCA) representation of cuticular compounds of *Dolichoderus bispinosus* workers (black circles), *Juditha molpe* caterpillars (black triangles), *Nymphidium chione* caterpillars (orange triangles), and *Pheidole biconstricta* workers (orange circles).
