## Supplementary table for "On the adaptive meaning of chemically insignificant cues in multitrophic caterpillar-ant-plant symbioses"

**Table S1.** Cuticular compounds that were found on riodinid caterpillars and associate ants (relative abundance % ± Standard Deviation). (-) = compound not detected.

| | | Caterpillar species | | Ant species | |
|---|---|---|---|---|---|
| Compound | Retention time | *Juditha molpe* | *Nymphidium chione* | *Dolichoderus bispinosus* | *Pheidole biconstricta* |
| (Z)-$C_{19}$ | 7.716 | - | - | 1.10 ± 1.38 | - |
| (Z)-$C_{21}$ | 10.873 | - | - | 0.11 ± 0.26 | - |
| $C_{21}$ | 11.375 | - | 3.41 ± 7.08 | 0.16 ± 0.27 | 0.05 ± 0.16 |
| Unidentified | 11.598 | - | - | - | 0.60 ± 1.92 |
| Ethyl linoleate | 11.966 | - | - | 0.12 ± 0.20 | - |
| Unidentified | 13.010 | - | - | - | 0.11 ± 0.26 |
| 3;7DiMe$C_{21}$ | 13.135 | - | - | 0.17 ± 0.31 | - |
| (Z)-$C_{23}$ | 14.104 | 0.82 ± 1.65 | 2.74 ± 2.71 | 4.76 ± 3.99 | 9.47 ± 8.75 |
| $C_{23}$ | 14.607 | - | - | 0.18 ± 0.32 | 1.01 ± 1.00 |
| 11-Me-$C_{23}$ | 15.183 | 1.2 ± 2.4 | 5.80 ± 1.65 | 0.74 ± 0.52 | 0.34 ± 0.48 |
| $C_{24}$ | 16.175 | - | - | - | 0.14 ± 0.31 |
| 3;7DiMe$C_{23}$ | 16.323 | - | - | 0.08 ± 0.27 | 0.31 ± 0.45 |
| (Z)-$C_{25}$ | 17.235 | - | - | - | 1.05 ± 1.07 |

| | | | | | |
|---|---|---|---|---|---|
| Unidentified | 17.346 | - | - | 9.86 ± 6.07 | 11.01 ± 9.10 |
| $C_{25}$ | 17.695 | 11.01 ± 1.77 | 33.40 ± 6.53 | 1.03 ± 0.88 | 2.25 ± 1.71 |
| 13;11-Me-$C_{25}$ | 18.226 | 0.49 ± 0.99 | - | - | 0.03 ± 0.11 |
| 5-Me-$C_{25}$ | 18.336 | - | - | 0.05 ± 0.18 | 0.10 ± 0.23 |
| Unidentified | 18.462 | - | - | 0.04 ± 0.12 | - |
| 3-Me-$C_{25}$ | 18.788 | - | - | - | 0.33 ± 0.54 |
| Unidentified | 18.847 | 2.74 ± 3.97 | - | 3.64 ± 2.16 | 0.34 ± 0.41 |
| $C_{26}$ | 19.176 | 0.14 ± 0.29 | 7.62 ± 2.96 | - | 0.04 ± 0.15 |
| 3;7-DiMe-$C_{25}$ | 19.355 | 0.35 ± 0.70 | - | 0.53 ± 0.51 | - |
| 14;12-Me$C_{26}$ | 19.670 | - | - | - | 16.65 ± 11.99 |
| (Z)-$C_{27}$ (2) | 20.198 | - | - | - | 9.83 ± 8.07 |
| $C_{27}$ | 20.305 | 16.25 ± 3.03 | 12.61 ± 2.14 | 26.19 ± 12.58 | 5.17 ± 3.20 |
| 3-Me-$C_{27}$ | 20.605 | 8.14 ± 9.58 | - | 8.72 ± 6.08 | 3.86 ± 1.28 |
| 13;11-Me-$C_{27}$ | 21.073 | 0.74 ± 1.48 | - | 1.79 ± 1.33 | - |
| Unidentified | 21.128 | 0.22 ± 0.44 | - | 0.14 ± 0.19 | - |
| 7-Me-$C_{27}$ | 21.209 | 0.27 ± 0.55 | - | 1.09 ± 0.88 | 0.09 ± 0.31 |
| 5-Me-$C_{27}$ | 21.345 | 0.47 ± 0.94 | - | 0.42 ± 0.51 | - |
| Unidentified | 21.497 | 2.59 ± 3.69 | 14.56 ± 15.28 | 2.12 ± 0.99 | 0.27 ± 0.46 |

| | | | | | |
|---|---|---|---|---|---|
| 3-Me-$C_{27}$ | 21.647 | - | - | 0.11 ± 0.28 | - |
| 5;9-DiMe-$C_{27}$ | 21.804 | 0.26 ± 0.53 | - | - | - |
| $C_{28}$ | 21.993 | 0.24 ± 0.48 | - | 0.66 ± 0.49 | - |
| Unidentified | 22.085 | - | - | 0.09 ± 0.21 | 0.03 ± 0.10 |
| 3;7-DiMe$C_{27}$ | 22.194 | - | - | 0.08 ± 0.18 | 5.87 ± 8.26 |
| 16;14;12-Me-$C_{28}$ | 22.441 | - | - | - | 0.20 ± 0.36 |
| $C_{29}$ | 22.865 | 31.76 ± 11.48 | 12.11 ± 3.47 | 9.6 ± 5.32 | 3.05 ± 4.80 |
| 13;15-Me-$C_{29}$ | 22.962 | 1.78 ± 2.06 | - | 2.62 ± 1.32 | 5.41 ± 6.14 |
| 5;3-Me-$C_{29}$ | 23.078 | 0.20 ± 0.41 | - | 1.66 ± 1.04 | - |
| 15;12;9-Me-$C_{30}$ | 23.344 | - | - | 0.18 ± 0.38 | 0.11 ± 0.34 |
| 13-Me-$C_{29}$ | 23.762 | - | - | 1.5 ± 3.10 | - |
| 5-Me-$C_{29}$ | 24.023 | 0.24 ± 0.49 | - | 0.40 ± 0.38 | - |
| 11;15;13;17-Me-$C_{29}$ | 24.131 | 1.64 ± 2.45 | 3.98 ± 2.48 | 0.84 ± 1.55 | 0.15 ± 0.34 |
| 14-Me-$C_{30}$ | 24.316 | - | - | 0.19 ± 0.41 | 0.55 ± 0.90 |
| 13;17;15-DiMe$C_{29}$ | 24.428 | - | - | 0.04 ± 0.12 | - |
| $C_{30}$ | 24.639 | - | - | 1.87 ± 3.30 | - |
| 12;14;16-Me-$C_{30}$ | 25.039 | - | - | 0.97 ± 1.62 | 3.72 ± 5.47 |
| (Z)-$C_{31}$ (2) | 25.182 | - | - | 0.04 ± 0.12 | - |

| | | | | | |
|---|---|---|---|---|---|
| Unidentified | 25.459 | - | - | 0.49 ± 1.37 | - |
| (Z)-$C_{31}$ | 25.574 | 18.38 ± 6.42 | 3.42 ± 2.12 | 4.75 ± 5.74 | 2.32 ± 4.54 |
| 15-Me-$C_{31}$ | 25.677 | - | - | 1.86 ± 3.27 | 10.82 ± 15.79 |
| Unidentified | 25.815 | - | - | 1.60 ± 1.67 | - |
| 17;15-DiMe-$C_{31}$ | 25.909 | - | - | 0.23 ± 0.74 | 1.40 ± 2.33 |
| 15;13-Me-$C_{31}$ | 26.304 | - | - | 0.80 ± 1.55 | - |
| 5-Me-$C_{31}$ | 26.564 | - | - | 0.39 ± 0.37 | - |
| $C_{31}$ | 26.647 | - | 0.30 ± 0.90 | 0.10 ± 0.32 | - |
| Unidentified | 26.820 | - | - | 0.19 ± 0.42 | 0.24 ± 0.52 |
| 15;17-DiMe$C_{31}$ | 26.974 | - | - | 0.03 ± 0.12 | - |
| Unidentified | 27.232 | - | - | 1 ± 2.13 | - |
| 12;14-Me-$C_{32}$ | 27.673 | - | - | 0.33 ± 0.74 | - |
| $C_{33}$ | 28.024 | - | - | 0.69 ± 0.73 | 0.07 ± 0.23 |
| 17;15;13-Me-$C_{33}$ | 28.152 | - | - | 1.11 ± 1.27 | 2.85 ± 4.78 |
| Unidentified | 28.375 | - | - | 0.04 ± 0.14 | - |
| Unidentified | 28.786 | - | - | 0.30 ± 0.65 | - |
| Unidentified | 29.322 | - | - | 0.12 ± 0.38 | - |
| 5;3-Me-$C_{33}$ | 29.531 | - | - | 0.29 ± 0.50 | - |

| | | | | | |
|---|---|---|---|---|---|
| Unidentified | 29.701 | - | - | 0.05 ± 0.16 | - |
| 17;15;13-Me-$C_{35}$ | 29.790 | - | - | 1.35 ± 1.80 | - |
| 5-Me-$C_{35}$ | 30.277 | - | - | 0.15 ± 0.32 | - |
